# Potential impact of TE-derived sRNA on gene regulation in the grass *Brachypodium distachyon*

**DOI:** 10.1101/2022.04.05.487121

**Authors:** Michele Wyler, Bettina Keller, Anne C Roulin

## Abstract

In plants, the transcriptional and post-transcriptional repression of TEs involved the production of small interfering RNAs (siRNAs) that guide the *de novo* methylation and transcriptional silencing of TEs. Because siRNAs act via sequence complementarity, they indiscriminately target TEs and possible non-TE transcripts. TEs and their derived siRNAs might thus provide a reservoir for long-distance gene regulation. To test this hypothesis, we infected *Brachypodium distachyon* with *Mangnaporte oryzae*, the agent of Rice Blast. The infection caused the differential expression of multiple TE copies as well as a global change in gene expression. By sequencing the small RNA fraction, we identified TE-derived sRNA that are identical in sequence to motifs contained in the UTR region of differentially expressed genes. Our study opens new avenues of research to further investigate to what extent TEs may serve as a source for in trans gene regulation.

## Introduction

TEs are mobile DNA sequences that can increase their copy number and move from one location to another within the genome (Wicker, Francois Sabot, et al. 2007). Because of their transposition dynamics, TEs constitute a significant fraction of eukaryotic genomes (Wicker, Francois Sabot, et al. 2007; Tenaillon et al. 2010), hereafter referred to as the mobilome. TEs are also known to play an important functional role. Indeed, TEs capable of replication carry promoter sequences and open reading frames encoding the transposition machinery. In the simplest case, the insertion of such sequences across the genome interrupts functional elements (Bhattacharyya et al. 1990). More subtle effects at the regulatory level include the creation of alternative or new promoters, the alteration of epigenetic landscapes and the subsequent fine tuning of gene expression (Lisch 2013; Bourque et al. 2018; Qiu and Köhler 2020).

In the field of molecular ecology, TEs have attracted much attention for their effects on the expression of nearby genes and resulting phenotypic variation (Butelli et al. 2012; Lisch 2013; Wang et al. 2013; Hof et al. 2016; Thieme et al. 2017). However, TEs can also trans-regulate genes. For instance, the presence of repetitive elements in the genome can influence the 3D conformation of chromosomes (Grob and Grossniklaus 2019), which correlates with transcriptional activity (Rosin et al. 2008). TEs can also be disruptive in various ways and are consequently under tight control (Hollister and Gaut 2009; Lisch 2009; Ito et al. 2011; Lisch and Bennetzen 2011; Bucher et al. 2012; Fedoroff 2012). In plants, the transcriptional repression of TEs is mediated by the RNA-directed DNA methylation (RdDM) pathway (Matzke and Mosher 2014; Sigman and Slotkin 2016) and involves the RNA polymerase enzyme Pol IV. TE transcripts generated by Pol IV are copied to make double-stranded RNAs (dsRNAs), which are then processed into 24-nt small interfering RNAs (siRNAs) that guide *de novo* methylation and transcriptional silencing of TEs. Because siRNAs act via sequence complementarity, they indiscriminately target TEs and possible non-TE transcripts (McCue et al. 2012; A D McCue et al. 2013; Cho 2018). TEs and their derived siRNAs might thus provide a reservoir for long-distance gene regulation, which has been hypothesized and experimentally validated in *A. thaliana* for a few genes (McCue et al. 2012; A D McCue et al. 2013). However, how this process may further help individuals respond to stress and adapt to challenging environmental conditions is yet unexplored.

In this regard, retrotransposons are of special interest among TEs. These elements, which are particularly abundant in plant genomes (Vitte et al. 2014), are structurally and functionally related to retroviruses. Transposing through a copy-paste mechanism, retrotransposon RNA transcripts are reverse transcribed into double-stranded DNA that is then integrated into the genome to form new copies (Sabot and Schulman 2006). Because reverse transcription is inherently error-prone and mutagenic, retrotransposition events result in the emergence of retrotransposon populations with many variants (Josep M. Casacuberta, Samantha Vernhettes and Grandbastien 1997). Whether this mechanism helps retrotransposons adapt to the genome they populate by fostering diversification is under debate but it undoubtedly provides a substrate for siRNA diversity and increased gene regulatory potential upon exposure to environmental stress. This mechanism may therefore be crucial to local adaptation.

Here, we investigated to what extent TEs might provide a reservoir for sRNA production and long-distance gene expression regulation. To do this, we used the model grass *Brachypodium distachyon* and the fungus *Mangnaporte oryzae* as a stress. *Mangnaporte oryzae* is the causal agent of the rice blast disease (Routledge et al. 2004; Dean et al. 2005). Rice blast can affect a wide range of temperate grasses and cereals, among which *B. distachyon* (Routledge et al. 2004). Using a genome wide RNA-Seq and sRNA-Seq approach, we investigated gene, sRNA and TEs expression pattern during rice blast infection. We identified differentially expressed TE-derived sRNA that are identical in sequence to motifs contained in the UTR region of differentially expressed genes. We thus provide a list of potentially *trans*-regulated genes for further molecular validation.

## Results

### TE annotation

To investigate the transcriptional effect of TEs on sRNA production, we only used elements annotated with high confidence. The retrotranspon annotation was recently updated by (Stritt et al. 2019) and included 4,612 TEs from 40 different families. We further reannotated the position of DNA transposons in the genome. We were able to identify 31,613 high confidence DNA TE copies that cluster into 144 TE families among which 30 had not been described previously. All the TE consensus sequences used in this study have been submitted to the publicly available database TREP (Wicker, François Sabot, et al. 2007, https://botserv2.uzh.ch/kelldata/trep-db/index.html).

### Differentially expressed genes and TEs

To investigate the transcriptional impact of a fungal infection, we performed RNA-seq of the reference *B. distachyon* accession Bd21 infected with *Magnaporthe oryzae* after 24 and 96 hours, with three replicates for each condition. On average 27’438’700 reads were mapped to the genome (94%). We then performed a differentially expression analysis comparing the control plants with the two infection time points. After 24 hours, only seven out of the 23’181 genes were differentially expressed in Bd21. After 96 hours, however, 3’262 genes were significantly differentially expressed compared to the control (1’607 down, 1’655 up regulated genes, Fig. 1A).

**Figure 1.**
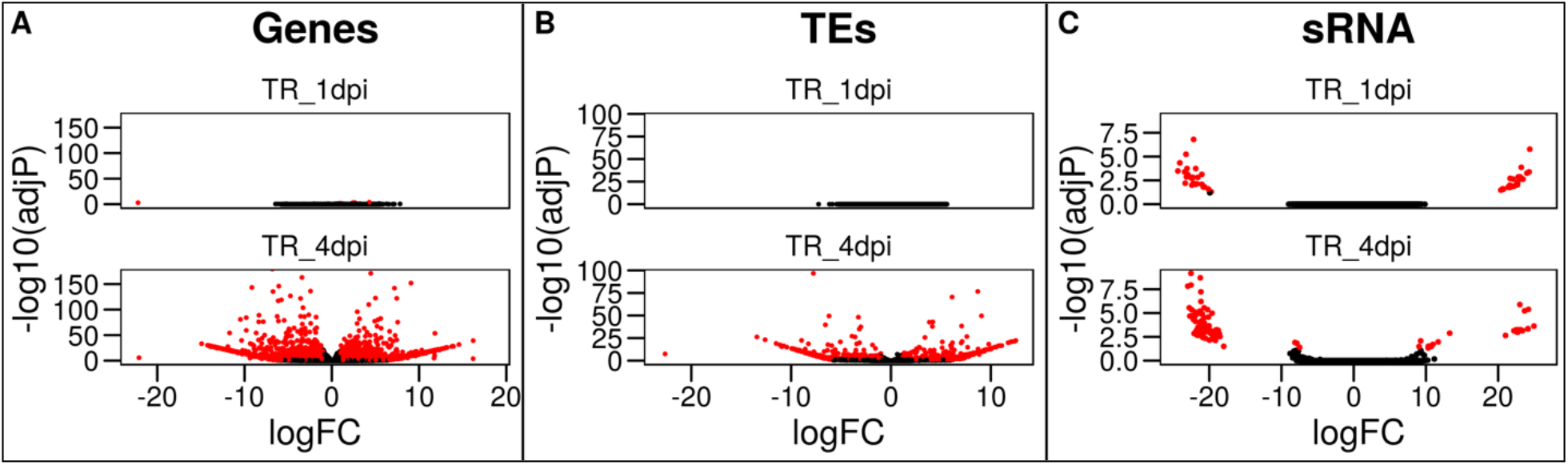
Volcano Plot for genes (A), TEs (B) and sRNA (C). Significantly differentially transcribed elements (absolute log Fold Change > 1, multiple testing adjusted p values < 0.05), are colored in red, the remaining in black. The upper plots are representing the expression after 24 hours post infection. The bottom plots depict the transcription after 96 hours.

We performed the same analyses for TEs. As observed for genes, none of the 31’631 annotated TEs were differentially expressed after 24 hours post infection in Bd21. On the other hand, we identified 930 differentially expressed TE copies after 96 hours compared to the control (327 down, 603 up regulated TEs, Fig. 1B).

We used a randomForest classifier to assess which genomic features affect the expression of TEs upon *M. oryzae* infection. The model had a predictive accuracy of 84% (Fig. 2A) and identified five most important variables (in order): the number of methylable sites of cytosine in a CHG context (where H can be C, A, T), the number of methylable of cytosine in a CG context, the length of the element, the number of methylable of cytosine in a CHH context and the GC content. We also found that differentially expressed TEs are significantly longer and and display a lower GC content in all three contexts (*p* < 0.05, Fig. 2B).

**Figure 2.**
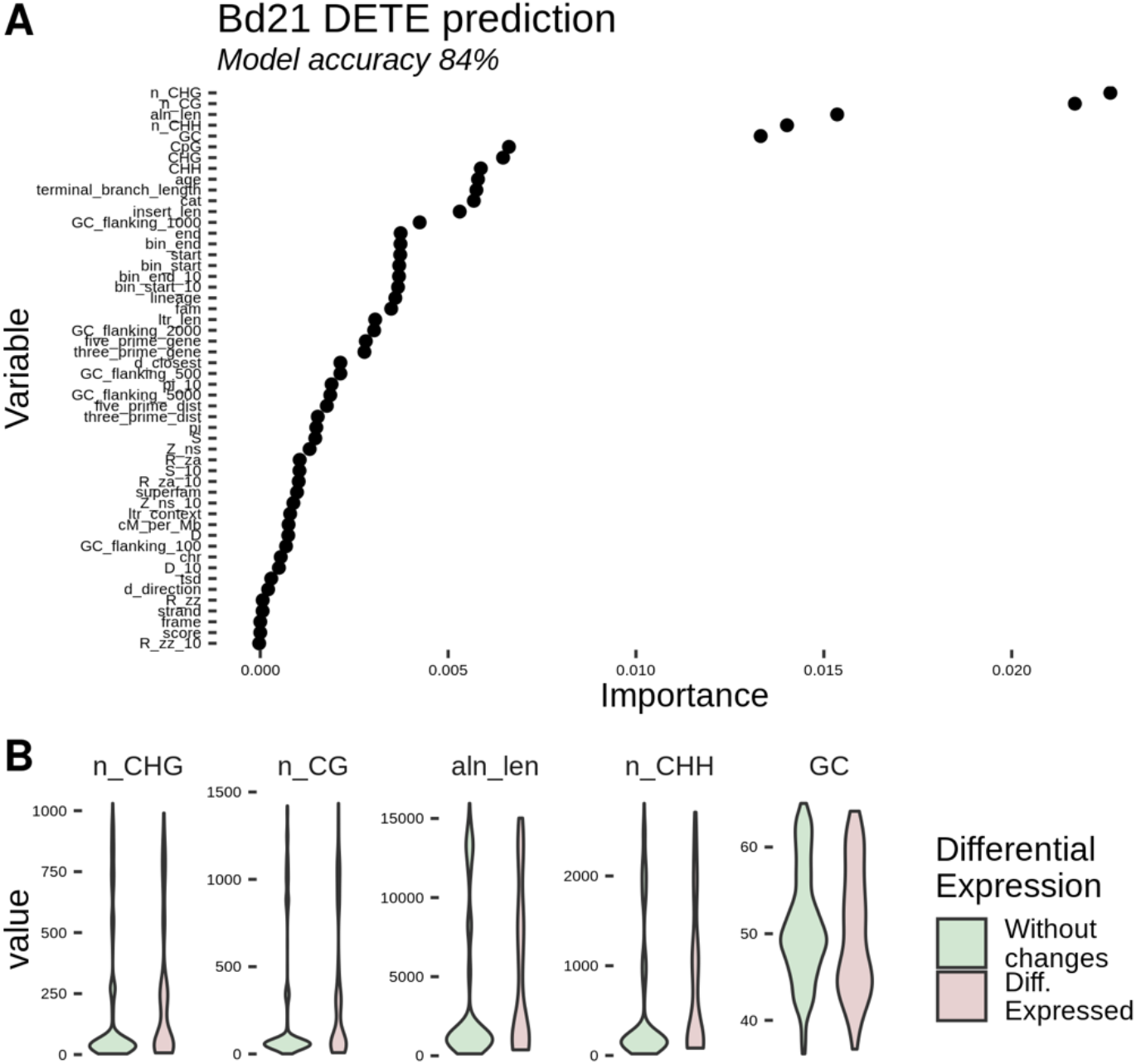
Random Forest classification. A) Variable importance for differentially transcribed TEs as ranked by the random forest classifier. B) Comparison of the distribution of the five most important classifier between TEs with (red) or without (green) differential expression. n_CHG is the amount of Cytosines in a CHG context in the TE (where H can be C, T or A). n_CG is the amount of Cytosines in a CG context in the TE. aln len is the length of the annotated TE copy. n_CHH is the amount of Cytosines in a CHH context in the TE. GC is the Guanine/Cytosine ratio of the TE copies. All five traits differ significantly (p value < 0.05).

We also sequenced the small RNA fraction under control and infected conditions and performed subsequently a sRNA expression analyses. In contrast to what we observed for genes and TEs, we identified 50 sRNA sequences already differentially expressed after 24 hours compared to the control (25 down-, 25 upregulated). Similarly, after 96 hours, we counted 114 sRNA sequences whose expression was significantly different compared to the control (84 down-, 30 upregulated).

### sRNA target site shared by genes and TEs

Silencing mechanisms rely on sequence homology between sRNA and the target sequence. To investigate the possibility of trans regulation between TEs and genes, we defined two possible scenarios (Fig. 3). First, a TE is silenced and the gene displaying sequence similarity is therefore expressed. Alternatively, the increased expression of a TE could lead to a higher production of sRNA which may silence gene expression through sequence complementarity. Applying stringent filtering criteria (see methods), we identified four up-regulated genes and three down-regulated genes (Table 1) potentially regulated by TE-derived sRNA. Interestingly, all seven candidate genes harbored sequence complementary to the sRNA sequences in their UTR regions (Fig. 4). These regions are known regulators of gene expression.

**Figure 3.**
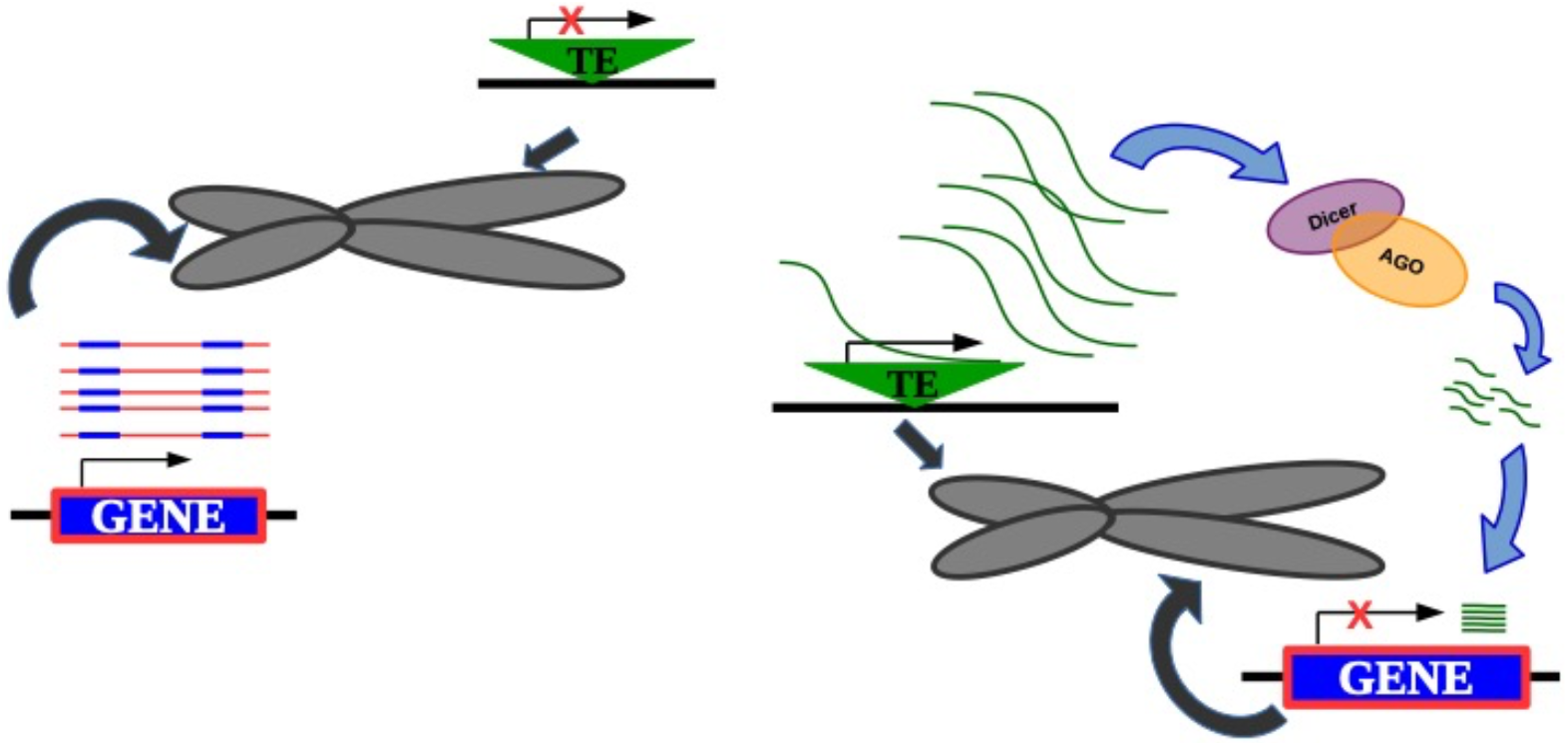
Transcription scenarios. A) TEs are silenced and genes are transcribed. B) TEs are transcribed, the resulting RNA is processed into sRNA that cause a reduction in gene expression through trans-regulation.

**Figure 4.**
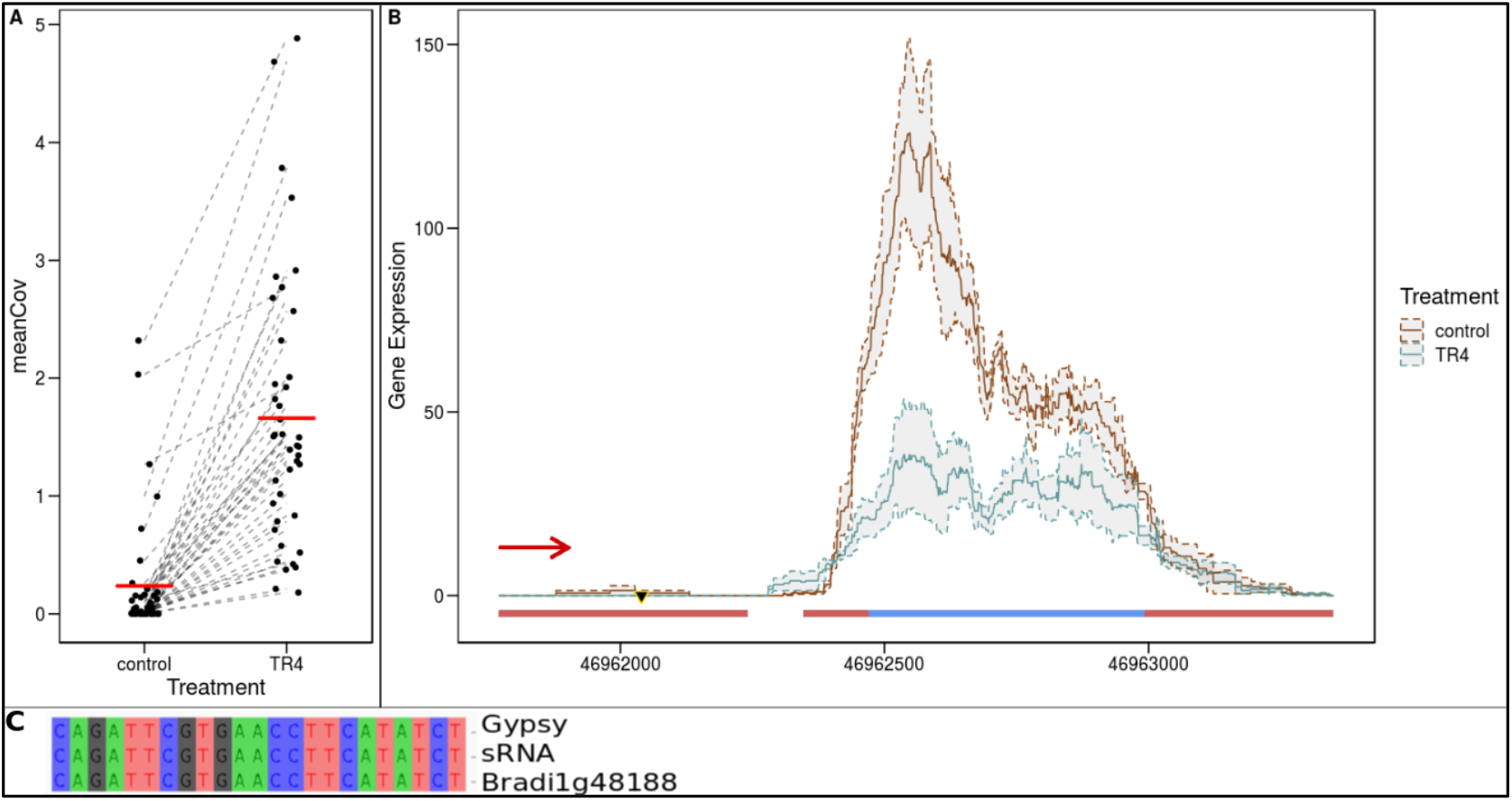
RNA coverage of A. Gypsy Copies with a shared sequence with B. Bradi1g48188. C. Alignment of shared sequences between Gypsy, sRNA and Bradi1g48188. Blue bars indicate exons, red bars UTRs. The sRNA sequence is indicated using the black triangle. The transcription direction is indicated by the arrow.

**Table 1.** Candidate genes regulated by TEs in Bd21 after M. oryzae infection.

| Gene ID | TE Family | sRNA | Domains (Pfam) | Gene Expression |
| --- | --- | --- | --- | --- |
|  |  | Length (bp) |  |  |
| Bradi1g68200.1 | Copia BdisC242 | 16 | DUF1618 | down |
| Bradi1g43423.1 | Gypsy (Multiple) | 24 | Retrotrans_gag | down |
| Bradi1g48188.1 | Gypsy (Multiple) | 23 | - | down |
| Bradi3g05470.2 | Copia BdisC010 | 24 | NB-ARC, Rx_N, LRR, AAA, TniB | up |
| Bradi2g47520.2 | Copia BdisC010 | 24 | AWPM-19-like family protein | up |
| Bradi2g59590.1 | Copia BdisC010 | 24, 22, 21 | - | up |
| Bradi2g29278.1 | Gypsy (Multiple) | 21 | - | up |

As an example, multiple TE copies from various Gypsy families shared an exact sequence with the UTR region of Bradi1g48188 (Fig. 4). The infection with *M. oryza* resulted in a significant increase in transcription of these copies (*p* < 0.05, Fig. 4A). We also observed a significant higher abundance of sRNAs putatively derived from these Gypsy elements and a significantly reduced expression of Bradi1g48188 upon infection (Fig. 4B, *p* < 0.05). To characterize the candidate genes, we searched for gene homologs described in other plant species but also scanned for known protein domains (Table 1). We found no known homologs except for Bradi3g05470.2 which is a homolog of the *Arabidopsis thaliana* gene AT3G46710.1 and contains well-described resistance gene domains such as NB-ARC and LRR (Urbach & Ausubel, 2017; Wang *et al.*, 2020). Two other candidate genes have known stress related domains. The AWPM-19-like family domain of Bradi2g47520 was shown to be related with stress tolerance through abscisic acid (ABA) regulation in rice (Chen *et al.*, 2015). Furthermore, the DUF1618 domain of Bradi1g68200 has unknown function, however this monocot specific domain was previously shown to be involved in stress response and fitness of rice (Wang *et al.*, 2014). One gene (Bradi1g43423.1) contains a retrotrans_gag domain, a typical domain present in TEs.

## Discussion

In this preliminary pilot study, we investigated the interplay between gene expression, TE activity and sRNA production in *B. distachyon* during an infection caused by the rice blast agent *Magnaporthe oryzae*. We identified a large number of differentially expressed genes and TEs over the infection time. This overall pattern reflects the rewiring of gene expression as a response to the pathogen attack. Furthermore, we identified a putative link between TE-derived sRNA production and gene expression levels regulation for a small number of candidates, indicating that TEs may have a limited a role on long-distance gene regulation.

TEs may play a fundamental role in stress responses (*e.g.* Makarevitch et al. 2015), however, transcriptomic approaches are often used to infer stress responsive genes (Chen et al. 2017; Powell et al. 2017). Here, we investigated the reaction of *B. distachyon* upon infection by the rice blast agent *M. oryzae*. We also performed a differential expression analysis for TEs. We found that the regulation pattern of TEs over time resembles the pattern visible for genes and thus hypothesized that the rice blast infection causes a genome wide deregulation of both genes and TEs. The observed upregulation of the transcription of specific TEs under stress conditions matches previously described patterns observed in various organisms under different stresses (Fouché et al. 2020; Liang et al. 2021) and supports the hypothesis of a role of TEs in genomic and/or transcriptomic innovation (McClintock 1947). However, our experiment reveals also a similar number of TEs that are significantly down-regulated compared to the control condition.

Features that impact TEs activity include methylation levels, integrity and age (Stritt et al. 2019). Although part of this work focused on the prediction and annotation of DNA TEs, we excluded these elements to avoid technical difficulties. Specifically, estimating the age of single copies is more difficult for DNA transposons than for retrotransposons for which insertion age can be estimated for single copies using LTR sequence identity (SanMiguel et al. 1998). However, DNA transposons are characterized by short terminal inverted repeats (TIR, Wicker and Keller 2007). Pace and Feschotte (2007) published a method to estimate DNA transposon insertion date. The described method is yet applicable only to a limited number of families between close related species, and was therefore not suitable for our study. In particular MITEs are miniaturized non autonomous TEs and hence lacking features suitable for insertion dating (SanMiguel et al. 1998). Non autonomous elements are incomplete regarding their coding sequence and use the transcripts from intact elements to transpose (Roffler et al. 2015). We therefore restricted our Random Forest analysis to the extensively characterized retrotransposons from Stritt et al. (2019) to investigate the causative characteristics of the TEs expression upon *M. oryzae* infection.

We chose over 50 variables (*e.g.* TE length, age, CG content) to test whether specific genomic features may affect the differentially expression of each specific TE copy upon stress exposure. From all the variables we tested, the model prioritized a subset of five. Interestingly, all these features are related to the amount of cytosines contained in the TE copy. In fact, differentially transcribed TEs differ from the remaining TEs by being longer and containing more C nucleotides in the three different methylable context, causing a lower GC content. Although the number of methylable cytosines is classified as important by the model, the measured methylation level itself is considered less influential for the classifier. This observation may be explained by the fact that our experiment consider TE methylation levels under control conditions only. Actually, a number of previous works are indicating changes of methylation levels caused by the exposition of plants to stress (Bilichak et al. 2012; Qian et al. 2019; Rajkumar et al. 2020). Subsequently the silencing of TEs becomes permissive and lead to the transcription of TEs (McCue and Slotkin 2012). However, our findings are in line with (Catoni et al. 2017) which showed an increase of epiallelic switches in regions of the genome with higher cytosine content.

We specifically looked for genes that might be regulated by TE-derived sRNA. By doing so, we identified a set of candidate genes in the reference Bd21 that were differentially expressed upon infection and harbored TE-derived sRNA sequence in the regulatory sequences. Interestingly, we detected the presence of the sequences of interest in the annotated UTR regions, region known to play an important role in the transcriptional regulation of genes (Srivastava et al. 2018), for all candidate genes. Altogether, the fact that genes involved in defense response are potentially regulated by TE-derived sRNA that would target their regulatory sequences suggests that our analysis highlighted strong candidates for functional validation. Yet, we identified only a limited number of candidate genes putatively regulated by TE-derived sRNA. While we applied stringent filtering criteria, our results are in line with previous works performed in *A. thaliana* (e.g. Andrea D. McCue et al. 2013) and suggest that albeit limited, TE-derived sRNA may contribute to specific gene expression patterns in plants.

## Material and Methods

### Transposable element annotation

Different annotation method were applied depending on the class of the TEs. For retrotransposons, the manually curated annotation available from Stritt et al. (2019) was used. In this database, multiple descriptive characteristics like age, integrity or DNA methylation levels are available for each element beside the genomic coordinates. DNA transposons were annotated *de novo* in this project using a comparative genomic approach. Briefly, homologs in the *B. distachyon* accession Bd21 were identified using BLAST+ (Altschul et al. 1990). The upstream regions of each gene were extracted and aligned with Smith-Waterman local alignment algorithm available in the EMBOSS suite (Rice 2000). Mismatches between the two strands were subsequently blasted against the reference genome Bd21 and subsequently aligned. For each alignment, a candidate consensus sequence was formed and if the criteria defined by (Wicker, François Sabot, et al. 2007) were matched, assigned to a TEs family. Finally, the Bd21 genome was annotated with the newly identified families, together with the available TEs consensus sequences available in the TREP database (Wicker *et al.*, 2007, pp. https://botserv2.uzh.ch/kelldata/trep-db/index.html).

### Rice blast infectiont and plant material

Spores of rice blast *M. oryzae* genotype FR13 were cultivated on OMA plates (Hayashi et al. 2009) for 31 days at room temperature in the dark. Spores were removed from the plate and diluted in distilled water and 50% Tween20 (in 10 ml water) to reach a concentration of 200’000 spores per ml. The spore solutions were used to infect leafs of 15 days old *B. distachyon* genotypes Bd21 grown on 1/2 MS Medium. The plants were exposed to 16 hours light and 8 hours dark with 23°C and 40% humidity. Control plants were infected with a mock solution containing water and 50% Tween20. Control and infected plants were then incubated upside down on as described above for the growing conditions. After 24 and 96 hours, infected leaves were harvested and directly flash frozen. Total RNA was extracted using miRNeasy Mini kit (Qiagen) following the user manual and sRNA and RNA were sent to Novogene (Beijing) for sequencing.

### RNA processing

Raw RNA reads from all conditions were quality-checked using FASTQC (Andrews 2010). Raw RNA reads were mapped using STAR 2.5.3a (Dobin et al. 2013) with default settings on the *B. distachyon* reference genome Bd21 available from Phytozome (Goodstein et al. 2012). To obtain read counts for genes and TE, featureCounts 1.5.0 (Liao et al. 2014) was used allowing multi mapping reads (option -M) and using respectively the gene annotation available from from Phytozome (Goodstein et al. 2012) and the new TE annotation described above.

Differentially expressed genes and TE were obtained using DESeq2 1.22.2 (Love et al. 2014) embedded in RNAseqWrapper 0.99.1 (Schmid 2017) with a single factor and the different infection time points as a contrast. The analysis was performed only for gene or TE with at least five reads for all samples. Differentially expression was defined with log2FC above an absolute of 1 with an adjusted *p*-value (FDR) smaller than 0.05.

### sRNA processing

Removal of low quality bases, adapter trimming and length selection of the raw sRNA reads was performed using using fastp 0.19.6 (Chen et al. 2018). Only reads between 16 bp and 24 bp were kept (options *-length_required 16 -length_limit 24*). Mapping of sRNA reads was performed with ShortStack version 3.8.5 (Johnson et al. 2016) allowing multimappers (options *-nohp -bowtie_m 200*). To exclude cross kindom effects (Weiberg et al. 2013), all reads that map to the *Magnaporthe oryzae* genome reference MG8 (ensembl Fungi, Howe et al. 2020) were removed. Due to high repeatability of sRNA, read counting is based on the mapped sequence counts instead of an annotation of specific loci. Differentially expression was subsequently performed as described for mRNA.

### Identification of candidate genes

To identify TE-derived sRNA, reads from small RNA sequencing were mapped against the genomic TE sequences using *‘blastn-short’* (Altschul et al. 1990). Similarly, sRNAs were blasted against the reference transcript sequences available on Phytozome 12 (Goodstein et al. 2012) to identify the possible targets of specific sRNA. For both cases a threshold of 0 gaps, 0 mismatches and full-length hit was chosen.

To investigate the silencing capacity of TE in trans, genes, sRNA and TE were categorized into the two possible combinations: *i*) the rice blast infection leads the downregulation of TE and its derived sRNA and an overexpression of the gene harboring the corresponding target site for the TE-derived sRNA or *ii*) the infection causes the transcription of TE, causing a higher production of sRNA and consequently a downregulation of the gene harboring a target site for the TE-derived sRNA. Transcripts of the candidate genes were scanned using hmmscan from HMMER 3.1b (Finn et al. 2011) and the Pfam database (release 33.1, Bateman et al. 2004) to characterize functionally the genes of interest.

Methylation level of the candidate genes and TEs was assessed as described in (Wyler et al. 2020) using publicly available whole genome sequencing reads (PRJEB39651). Shortly, adaptor and low base quality reads were trimmed using trim_galore 0.4.5 (bioinformatics.babraham.ac.uk/projects/trim_galore) and subsequently mapped to the *B. distachyon* reference genome Bd21 using Bismark (Krueger and Andrews 2011) with Bowtie2 (Langmead and Salzberg 2012). After deduplication with deduplicate_bismark, the methylation level of cytosine was extracted using bismark_methylation_extractor (with options: *-comprehensive -bedGraph -CX-ignore 2 -ignore_r2 1*).

### TEs characterization

Previously published data (Stritt et al. 2019) was used to investigate the activity of differentially transcribed TEs towards *M. oryza* infection. The information is available for each single TE copy and comprise genetic, genomic and epigenetic data. To account for possible correlating traits, the random forest approach implemented in the the R package ranger (version 0.12, Breiman 2001) was used. The classifier algorithm was run using permutation and without replacement for 50,000 trees (options *importance = “permutation”, replace = F*) with the differentially transcription as a target variable. Subsequent testing of the most important predictor variables was performed using least square linear model and anova. Therefore, count data was square root transformed and percentages were arcsin-square root transformed.

## Data availability

All the raw sequencing data produced for the current study are available under the ENA project number PRJEB51973. Consensus sequences of newly discovered TE families are available from the TREP database (https://botserv2.uzh.ch/kelldata/trep-db/index.html).

